# Evaluating sampling strategies for the detection of avian influenza viruses in the environment

**DOI:** 10.1101/2025.09.18.677014

**Authors:** Allison K. Miller, Lia Heremia, Stephanie J. Waller, Sarah-Lou Blanchard, Jordan T. Taylor, Michelle Wille, Neil J. Gemmell, David Winter, Edwina J. Dowle, Jemma L. Geoghegan

**Affiliations:** Department of Anatomy, University of Otago, Dunedin, New Zealand; Department of Microbiology and Immunology, University of Otago, Dunedin, New Zealand; New Zealand Institute for Bioeconomy Science Limited, Lincoln, New Zealand; Centre for Pathogen Genomics, Department of Microbiology and Immunology, at the Peter Doherty Institute for Infection and Immunity, The University of Melbourne, Melbourne, Victoria, Australia; WHO Collaborating Centre for Reference and Research on Influenza, at the Peter Doherty Institute for Infection and Immunity, Melbourne, Victoria, Australia; New Zealand Institute for Public Health and Forensic Science (PHF Science), Porirua, New Zealand

**Keywords:** eRNA, sediment, water, ducks, flu, rt-qPCR, metatranscriptomics, RNA sequencing, virus

## Abstract

Highly pathogenic avian influenza (HPAI) viruses threaten humans, livestock and wildlife, highlighting the urgent need for early warning surveillance. Environmental RNA (eRNA) monitoring provides a safer, cost-effective and non-invasive alternative to direct pathogen testing, yet the effectiveness of different sample types for avian influenza viruses (AIVs) is unclear. We evaluated four eRNA-based sampling methods in urban waterfowl ponds over approximately one year. All methods combined with RT-qPCR and metagenomic sequencing detected AIV, but detections were asynchronous, likely reflecting low viral concentrations or detection limits. These results highlight both the promise and the current limitations of eRNA-based AIV surveillance.

**Graphic abstract:** 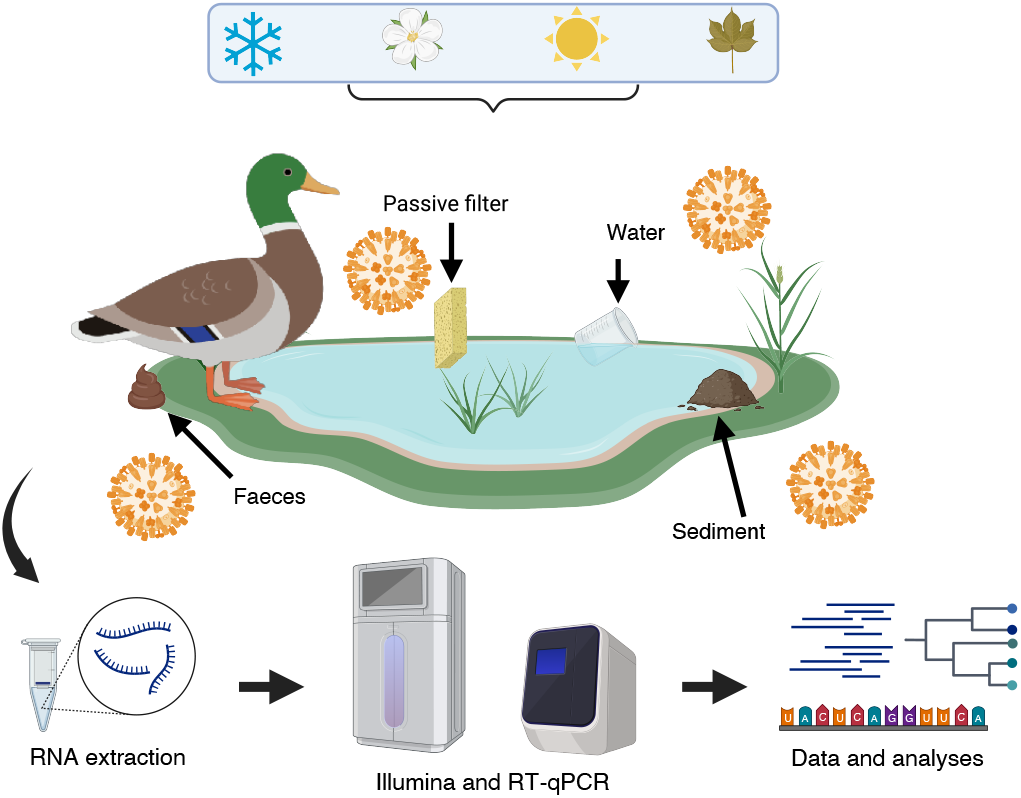

## Main Text

Avian influenza viruses (AIVs, species: *Alphainfluenzavirus influenzae*) are frequently detected in natural ecosystems, particularly among wild aquatic bird populations, where low pathogenic avian influenza (LPAI) viral strains typically circulate without causing overt signs of disease in their hosts (1). In contrast, the emergence, and global spread, of highly pathogenic avian influenza (HPAI) viruses – most notably subtype H5N1 – are a major concern for both animal and public health.Indeed, the increasing frequency of H5N1 spillovers from infected animals, such as poultry and other livestock, to humans, and the recent endemicity in cows (2), has heightened concerns about its potential to spark a global pandemic.

Efficient large-scale viral surveillance is essential for tracking viral emergence and incursion. Traditional surveillance approaches typically involve direct sampling from wild or domestic animals. While this method yields valuable insights into the virus’ host range, it is often time-consuming, expensive, requires specialised expertise and can expose fieldworkers to infection risks (3). In contrast, viral surveillance using environmental DNA (eDNA) or RNA (eRNA) relies on detecting viral genetic material shed into the environment during active infection. Sampling viral nucleic acids directly from the environment potentially offers an easier, safer, less invasive and more cost-effective alternative that can be implemented at scale (4) – an especially promising strategy for monitoring AIV (5).

Here, we assessed four eRNA sampling strategies and compared their sensitivity for detecting AIV using both metagenomics and reverse transcription quantitative polymerase chain reaction (RT-qPCR). From June 2024 to March 2025, we collected samples from two large urban ponds which are frequented by waterfowl, particularly introduced mallards (*Anas platyrhynchos*) in Dunedin Botanic Garden (DBG), Dunedin and Anderson Park (AP), Invercargill in Aotearoa New Zealand, ponds which are approximately 290 kilometres apart. From each, we collected four sample types, including environmental faecal samples, active filtered pond water, passive filtered pond water and sediment.

Ten fresh faecal samples were collected directly after observing ducks defecate, ensuring they were both fresh and attributable to waterfowl (predominantly mallards and other ducks), using sterile flock swabs placed into vials of DNA/RNA Shield (Zymo Research) during each monthly sampling. Water and sediment samples were collected from the same location at eight and six pond sites in DBG and AP, respectively. Water was filtered using Wilderlab (https://wilderlab.co/) Wetland kits and ∼30ml of submerged sediment was collected from ∼10 cm below the surface. Passive filters consisted of sterile sponges that were deployed on floats overnight for ∼16 hours (Supplementary Data 1). Pond water pH, salinity, dissolved oxygen and temperature data were collected via YSI ProPlus 2030 (Xylem Inc) and pH tester H198128 (Hanna) during each sampling trip (Supplementary Data 2). In addition, temperature and light Pendant loggers (Hobo) were submerged in ponds for the entire study (June 2024 to March 2025) (Supplementary Data 2).

Overall, we collected faecal samples (n = 213), sediment (n = 136), active filtered water (n = 132) and passive filtered water (n = 157). Total RNA was extracted from each sample separately using the ZymoBIOMICS MagBead RNA kit (Zymo Research) following a previously described protocol (6). Slight modifications were made for the water, sediment and passive filtered water samples: the DNA/RNA Shield was removed from the Wilderlab filter; sediment was well mixed with DNA/RNA Shield and then subsampled (800ul); and a corner of the passive sponge filter was subsampled into DNA/RNA Shield. Equal volumes of the extracted RNA were pooled into 80 sequencing libraries based on environmental sample type, collection date and location. The pooled samples were subject to total RNA sequencing using the Illumina NovaSeqX platform following ribosomal RNA depletion with Ribo-Zero-Plus (Illumina), generating an average of 94.5 million paired-end reads per library (range: 68.9 –154.1 million). In addition, the 80 pooled libraries were subject to RT-qPCR, targeting matrix protein 1 to confirm AIV metagenomic detection and sensitivity (Supplementary Data 3).

Sequencing reads were quality trimmed using Trimmomatic (v0.38) (7) and assembled *de novo* using MEGAHIT (v1.2.9) (8). Assembled contigs were annotated using a sequence similarity-based approach where contigs were annotated at the nucleotide level against a custom AIV database and further confirmed using the BLASTn webtool. Contig abundance was determined by mapping to sequencing reads using Bowtie2 (v2.5.4) (9) and read counts were standardised to reads per million (RPM).

AIV transcripts were identified across all sample types (Figure 1), although there was minimal overlap between sample type and sampling month. Notably, AIV detection via metagenomics was very limited during spring and early summer (October to January) possibly due to low viral concentration, which may reflect reduced duck activity in ponds during breeding season (10) or low AIV prevalence in the duck population. Surprisingly, AIV was detected in only a single faecal sample from AP using metagenomics, yet at a relatively high abundance of ∼35 RPM, substantially higher than in other environmental sample types (Figure 1). Although no AIV was detected in faecal environmental samples from DBG metagenomic data, AIV transcripts were present in all other sample types.

**Figure 1.**
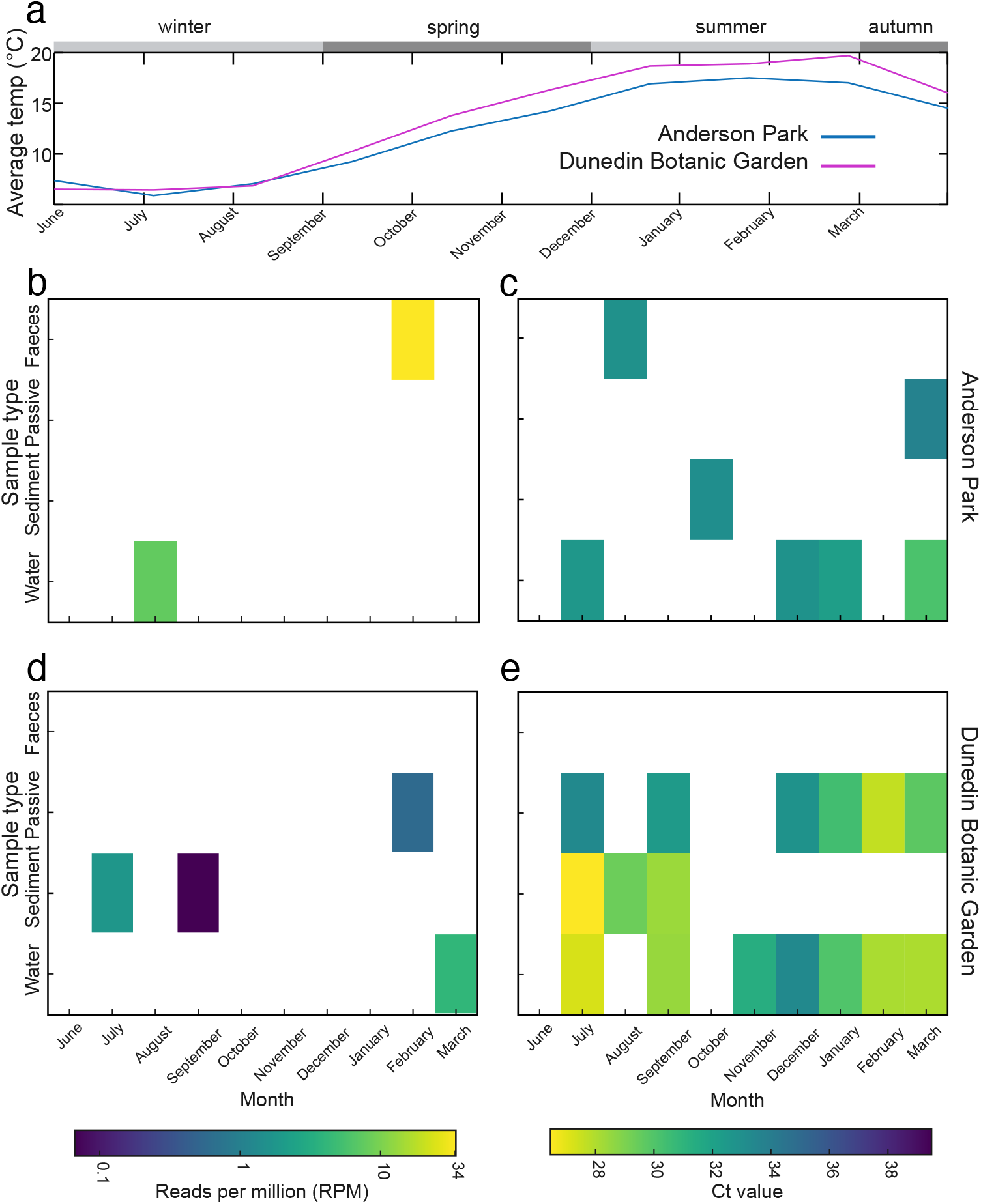
Occurrences of AIV detected by metagenomic analyses from June 2024 to March 2025. (**a**) Pond temperature (degrees Celsius) at the two sampling locations: Anderson Park (AP) in pink and Dunedin Botanic Garden (DBG) in blue. (**b & d**) Heatmap showing the abundance of AIV reads (per million) detected at each site (AP and DBG, respectively) for each month. (**c & e**) Heatmap showing the Ct value of detected AIV via RT-qPCR at each site (AP and DBG, respectively) for each month. In all heatmaps, a lighter colour (yellow) represents a higher viral load detected in a given sample.

We found no association between AIV abundance in metagenomic data (RPM) and Ct value using Spearman’s ρ correlation coefficient (*p* = 0.58 and *p* = 0.24 for AP and DBG, respectively). AIV detected via RT-qPCR was most frequently identified in active filtered water samples (55% of samples) compared to other sample types (35% of passive filtered water samples, 20% of sediment samples and 5% of faecal environmental samples) (Figure 1).

We found that all four sampling methods were able to detect AIV from waterfowl ponds; however, there was little consistency between sample types, detection method, locations and timing.Agreement between metagenomics and RT-qPCR detection was evaluated using McNemar’s test, which showed that qPCR detected significantly more AIV-positive samples than metagenomics (χ^2^ = 12.19, *p* = 0.0005). In addition, detection success varied by sample type for qPCR (χ^2^ = 13.36, *p* = 0.003), with filtered water yielding the highest AIV detection rate (55%), whereas metagenomics showed no significant difference among sample types (χ^2^ = 0.72, *p* = 0.87). The overall lack of consistency suggests environmental sampling coupled with metagenomic methods may produce frequent false negative results due to low concentrations of viral genetic material in eRNA. This inconsistencies among sample type may reflect the persistence of viral RNA in wetland environments after active infection has passed, in contrast to fresh faecal sampling which more directly reflects current infection status.

Despite being located 290km apart, both sampling locations exhibited similar and stable water parameters throughout the sampling period, except for temperature, which ranged from 3 – 24°C and peaked in February 2025 (Figure 1 and Supplementary Data 4). AIV transcripts were detected at both water temperature extremes (i.e. summer and winter) (Figure 1). Temperature, pH, and salinity are known to be important factors for the persistence of AIV in water, which can remain detectable for over 60 days at lower temperatures (11), and can even be subtype-specific (12).

While transcripts for all AIV genome segments were detected in our metagenomic data (Figure 2), transcripts for surface proteins hemagglutinin (HA) and neuraminidase (NA) were detected in faecal, sediment and passive filtered water samples only. These transcripts were first aligned using MAFFT (v7.526) (13) along with their closest genetic relatives obtained from NCBI GenBank and time-calibrated maximum likelihood phylogenetic trees were estimated using IQ-TREE (v3) (14) using Least Squares Dating. Phylogenetic analysis placed all environmental AIV detections within distinct Australasian clades (Figure 2). All HA and NA sequences clustered with those previously sampled from New Zealand mallards, except for N8 which was closely related to samples from South Australian waterbirds, most likely due to a lack of N8 samples from New Zealand. The phylogenetic position of HA and NA segments and the relatively large time gap between these sequences and the last sampled common ancestors, suggest these LPAI lineages are not new introductions to New Zealand.

**Figure 2.**
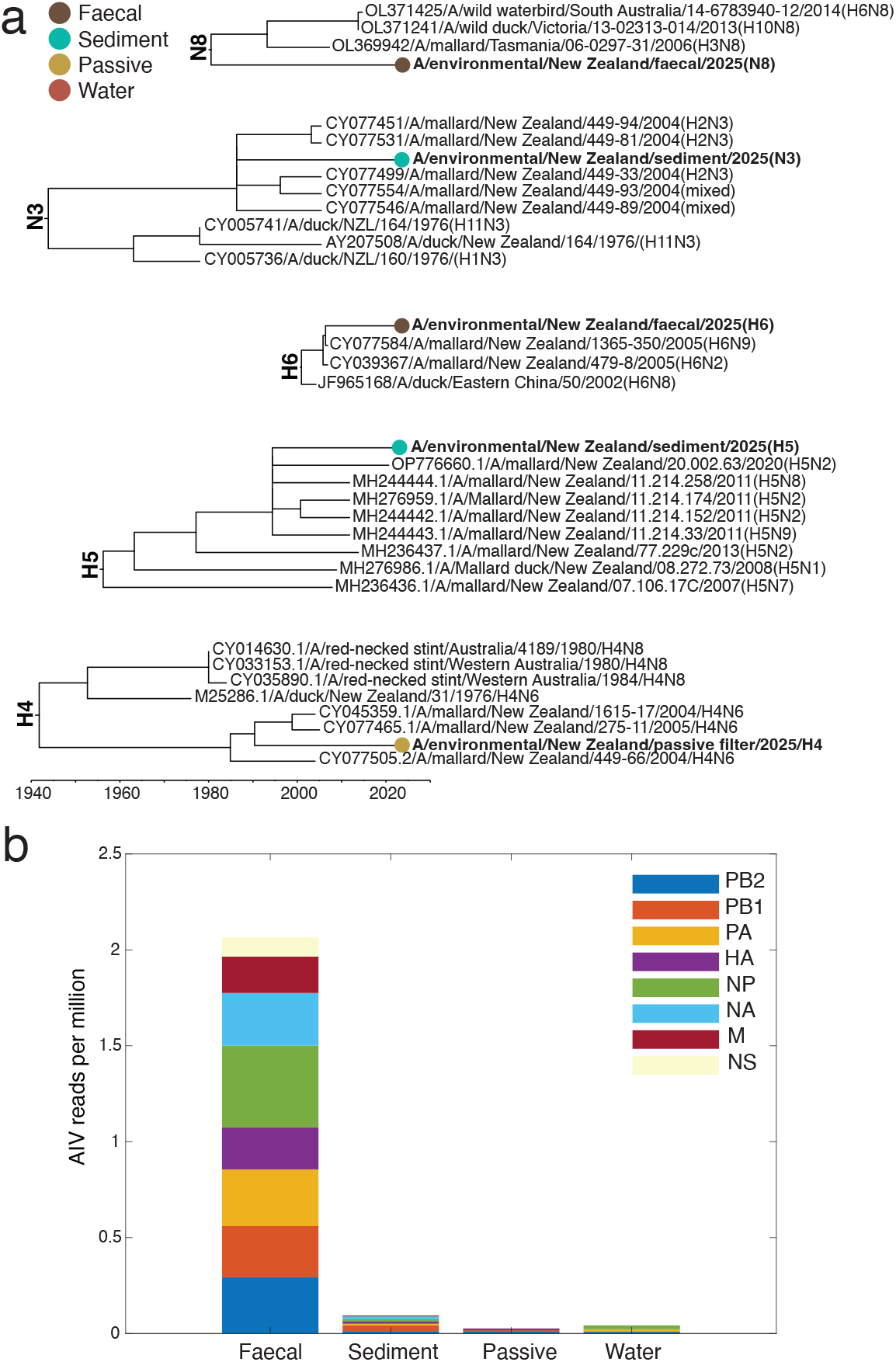
Phylogenetic and transcript results recovered from metagenomic analyses. (**a**) Time calibrated maximum likelihood phylogenetic analyses of the surface protein (HA and NA) sequences with their closest genetic relatives from NCBI GenBank. Colours correspond to the four sample types (faecal, sediment, passive water filters and active water filters). (**b**) Bar plot depicting the AIV segments reads (per million) recovered in each sample type. Colours correspond to the eight AIV segments (PB2 = Polymerase basic protein 2, PB1 = Polymerase basic protein 1, PA = Polymerase acidic protein, HA = Hemagglutinin, NP = Nucleoprotein, NA = Neuraminidase, M = Matrix protein 1 and NS = Non-structural protein 1).

To date, HPAI H5N1 has not been detected in New Zealand. However, LPAI viruses have been frequently identified in waterfowl and other aquatic birds through active surveillance efforts (15). This study represents an important first step in assessing the environmental detectability of AIV and evaluating the effectiveness of different environmental sampling approaches over time. Because viral RNA can persist in wetland environments, exact infection timing is not accurately captured, and HA-NA combinations cannot be resolved, from non-faecal environmental sampling. Nevertheless, given the immediate threat HPAI poses to public health, agriculture, domestic animals and wildlife, there is an urgent need to expand and refine current surveillance strategies. Enhanced environmental surveillance has the potential to serve as an early warning system for viral incursions and will be critical in supporting efforts to mitigate the risk of HPAI introduction.

## Supplementary information

**Supplementary Data 1**. Passive filter setup. Sterile sponges were attached to floats (**A**), deployed in the ponds (**B**), and left overnight for ∼16hrs. The sponges were then collected into sterile bags (**C**) and transported on ice to the laboratory where they were placed in a −80°C freezer until they could be extracted.

**Supplementary Data 2**. Environmental pond water parameters measured for this study from YSI (Xylem Inc YSI ProPlus 2030) and pH (Hanna tester H198128) meters (top), and Hobo Pendant Logger (averages, bottom).

**Supplementary Data 3**. Detailed methods of RT-qPCR for AIV matrix protein 1 detection.

**Supplementary Data 4**. Full Hobo Pendant Logger dataset collected from Anderson Park and Dunedin Botanic Gardens over the study period.

## Author contributions

Allison K. Miller: conceptualization, sampling, writing – original draft, review, and editing, investigation, methodology, validation, visualization, formal analysis, project administration, data curation. Lia Heremia: methodology. Sarah-Lou Blanchard: methodology. Jordan Taylor: methodology. Stephanie J. Waller: Sampling, writing – review and editing. Eddie Dowle: conceptualization, writing – review and editing. David Winter: writing – review and editing. Neil J. Gemmell: writing – review and editing. Jemma L. Geoghegan: conceptualization, investigation, funding acquisition, writing – original draft, review, and editing, analyses, project administration, supervision, resources.

## Ethics statement

No animals were handled for this research.

## Data availability statement

All sequencing data is available via NCBI’s GenBank database under accessions PX376089-PX376093.

## Acknowledgements

We would like to thank the Dunedin Botanic Garden, Anderson Park, and the Invercargill City Council teams, in particular Dylan Norfield, Derek Winwood for allowing us to collect samples from their grounds and to Eric Odell for showing AKM the DBG site. AKM would like to thank the Gemmell Lab members, particularly Jackson Treece, Sara Ferreira, and Joanne Gillum for their assistance, as well as JCC and PL for their sampling assistance. We appreciate the extra time spent by Rachel Hall and we thank Nicola McHugh and the University of Otago Zoology Department for letting us use their water monitoring equipment.

