## Supplementary figures and images for "Evaluating sampling strategies for the detection of avian influenza viruses in the environment"

### Supplementary Data 1

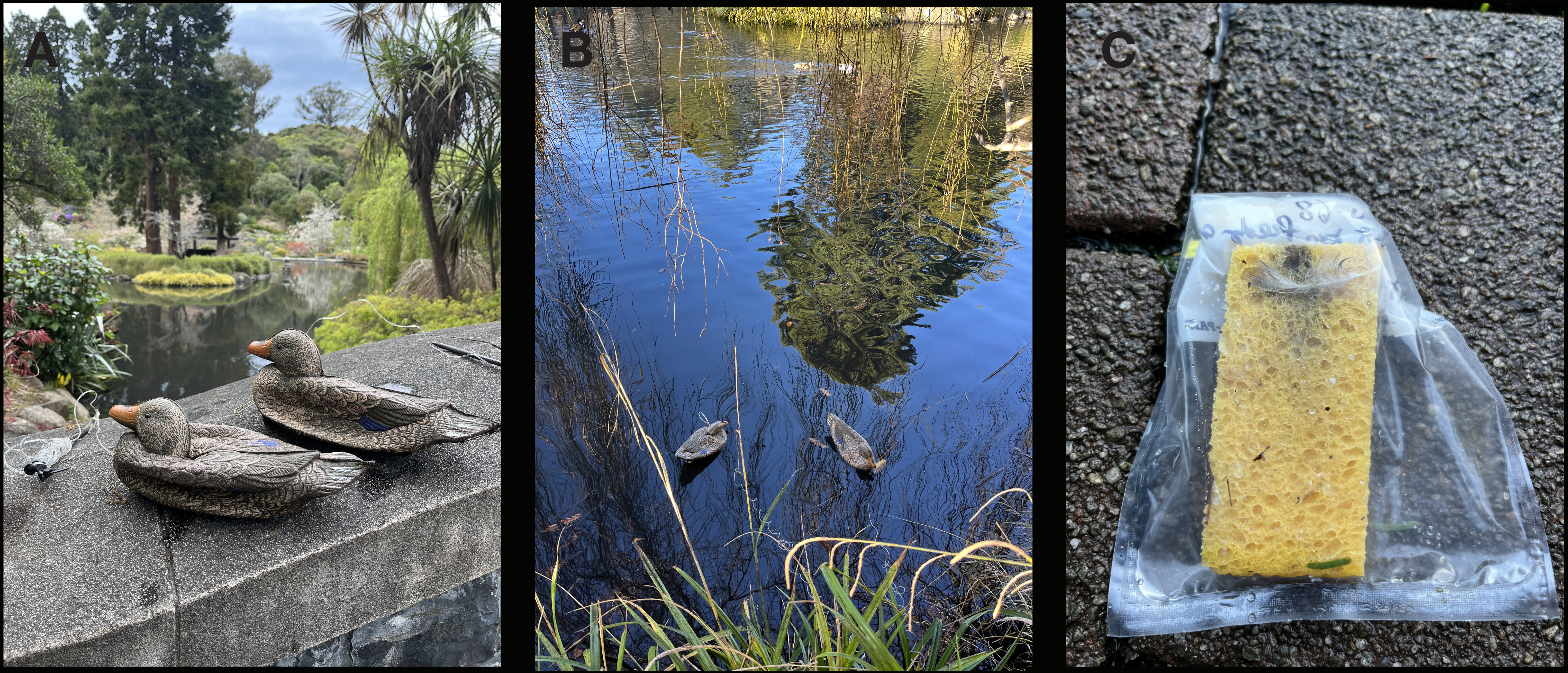
