## Supplementary Data 3 for "Evaluating sampling strategies for the detection of avian influenza viruses in the environment"

**RT-qPCR of Influenza Matrix Protein 1**

The 80 pooled libraries used for the total RNA sequencing were also subjected to RT-qPCR of the influenza matrix protein 1 (M1). The libraries were sent to Gnomix (<https://gnomix.com.au/>, Adelaid, Australia) where they were quality assessed and RT-qPCR assayed using Luna Universal One Step Rt-qPCR kits (E3005L) using the below primers and following the below protocols. Positive and negative controls were included and consisted of M1 from A/Turnstone/King Island/7164CP/2014(H6N8) (Wille et al. 2022) and off-target fur seal (*Arctocephalus forsteri*) viruses (those in the *Paramyxoviridae*). Rt-qPCR samples were performed in replicates of two. The positive control had a Ct value of ~14°C. Samples were considered positive for the presence of the M1 gene based on their Ct values and individual temperature profiles; Ct values of >35°C were excluded and samples were only considered positive if one of the two replicates were positive with a melting temperature (T_m_) of ~85°C (the same as the positive control).

### 1 Primers

**Primers from Fouchier et al. (2000):**

M52C: 5 -CTT CTA ACC GAG GTC GAA ACG-3

M253R: 5 -AGG GCA TTT TGG ACA AAG/T CGT CTA-3

### 2 Master Mix

| Reagent | Volume/Rxn (µl) | FINAL concentration |
| --- | --- | --- |
| Luna Universal One-Step Reaction Mix (2X) | 10 | 1X |
| Luna WarmStart® RT Enzyme Mix (20X) | 1 | 1X |
| Primer Forward working solution (10µM each) | 0.8 | 0.4µM |
| Primer Reverse working solution (10µM each) | 0.8 | 0.4µM |
| Nuclease-Free Water | 2.4 |  |
| Total | **15** |  |

### 3 Reaction Mix

| Reagent | Volume/Rxn (µl) |
| --- | --- |
| Luna In-Step Master Mix (see table 1) | 15 |
| RNA | 5 |
| Total | **20** |

### 4 Cycling conditions

#### 4.1 Luna One-Step

| Cycles | Temperature | time |
| --- | --- | --- |
| 1 | 55ºC | 10 min |
| 1 | 95ºC | 1 min |
| 45 | 95ºC  60ºC | 10 sec  30 sec* |

*Imaging
